# Brassinosteroid treatment reveals the importance of xyloglucan transglucosylase/hydrolase (XTH) genes in growth habit determination of twining common bean vines

**DOI:** 10.1101/2025.09.28.679089

**Authors:** Lena Hunt, Mariane S. Sousa-Baena, Angelique Acevedo, Leo Semana, Annabelle Wang, Rosemary A. E. Glos, Barbara A. Ambrose, Charles T. Anderson, Joyce Onyenedum

## Abstract

- Brassinosteroids impact the development of G-fibers —specialized cells that generate tension in plants. To explore the functional and genetic relationships between G-fibers and twining stems of common bean, we applied an active brassinosteroid and a brassinosteroid inhibitor to perturb G-fiber development and probed these phenotypes through gene expression and anatomical analyses.
- Brassinosteroid treatment generated phenotypes that affected the three key features of twining: elongation, circumnutation, and G-fiber development. We examined anatomical and biochemical changes in the G-fibers through cross-sections, macerations, and immunohistochemistry. RNA sequencing and differential gene expression analysis allowed us to identify unique gene expression patterns for each treatment.
- Brassinosteroid treatment led to significantly elongated internodes with disrupted circumnutation and long, thin-walled G-fibers. In contrast, inhibitor treatment produced short internodes with thick G-fibers. These phenotypes corresponded with significant differential expression of XTH genes, both at the onset of elongation and later, during G-layer deposition. Detection of xyloglucan epitopes in the G-layer, along with in situ hybridization, confirmed active xyloglucan remodeling after twining.
- Our results confirm the presence of xyloglucan in the G-layer of common bean, underscoring its importance in G-fiber function, and suggests a regulatory role for *XTH* genes in shaping the twining growth habit through modulation of cell wall properties.

## Introduction

When it comes to morphology, plants can be shapeshifters. Diverse growth forms occur among close relatives; for instance, the Fabaceae family includes trees, shrubs, and herbs, as well as climbers such as woody lianas and twining vines (Luizon Dias Leme et al., 2021; Stoffers, 1979). Remarkably, even a single species like *Phaseolus vulgaris* L. (common bean) can alternate between shrub-like and vining growth habits, depending on environmental conditions or selective breeding (Kelly, 2001). The evolution of a climbing growth habit has been hypothesized to be a key innovation that promotes greater species richness in lineages that acquire it, relative to non-climbing sister taxa (Gianoli, 2004, 2015). This advantage may stem from climbers’ ability to adapt their growth to available support (Ricklefs & Renner, 1994), and to fine-tune their development by allocating more biomass to transport tissues and leaves rather than self-supporting structures (Gartner, 1991; Wyka et al., 2013). Such flexibility allows climbers to exploit a variety of physical and light environments more effectively. Their rapid and adaptable growth provides a competitive edge over self-supporting plants and often negatively affects their hosts by outcompeting them for light and altering forest succession (Paul & Yavitt, 2011; Schnitzer, 2024; van der Heijden et al., 2013).

Among climbing plants are numerous species that are both economically valuable and ecologically important. Key crops such as grapes, peas, hops, beans, sweet potatoes, cucumbers, and squash are climbers. However, the same traits that make this growth habit advantageous can also give rise to pernicious structural parasites, including bindweed (Jacobs, 2007), dodder (Sandler & Ghantous, 2019), redvine (Elmore et al., 1989), and invasive kudzu (Lindgren et al., 2013). Despite the economic, evolutionary, and ecological success of climbing plants, the mechanisms underlying this growth habit remain underexplored. While model organisms such as *Arabidopsis thaliana* and *Populus* spp. have been instrumental in uncovering the biology of herbaceous and woody growth forms, neither system is well-suited for studying the cellular and developmental mechanisms unique to climbing plants. Here, we used the common bean, a plant with a well-annotated genome (Goodstein et al., 2012; *Phaseolus Vulgaris 2.1*, 2020), and growth-habit plasticity (Kelly, 2001), as an experimental system to study twining vines.

Twining vines, or “twiners”, represent the largest group of climbing plants, including more than half of neotropical climbing species (Sperotto et al., 2023). Twiners climb by wrapping their main stem around a support, without relying on accessory organs such as tendrils or aerial roots (Sperotto et al., 2020; Vaughn & Bowling, 2011). In twining varieties of common bean, the transition from shrub to vine growth habit occurs in three developmental stages as the plant matures. First, the upper internodes begin to elongate farther than the lower, shrubby internodes. Second, the shoot tip exhibits exaggerated circumnutation – rhythmic and increasingly pronounced circular movements, sweeping in wide arcs as it searches for a support structure. Finally, once contact is made and the stem coils around the support, gelatinous fibers (G-fibers)—specialized contractile fibers—form along the concave side of the coil, putatively helping the stem grip the support and maintain its posture (Onyenedum et al., 2025).

The contractile function of G-fibers arises as the innermost secondary cell wall layer (the gelatinous, or G-layer) (Clair et al., 2018; Mellerowicz & Gorshkova, 2012). (Sometimes referred to as a tertiary cell wall (Gorshkova et al., 2018). Unlike the lignified secondary cell wall, the G-layer is composed mostly of longitudinally-oriented crystalline cellulose in a matrix of pectic rhamnogalacturonan-I (RG-I), mannans, and arabinogalactan proteins (Bowling & Vaughn, 2008; Sugiyama et al., 1993). The very presence of xyloglucan in the G-fiber, however, remains a matter of debate in the literature, with early literature calling xyloglucan the most abundant non-cellulosic component (Baba et al., 2009, p. 2008; Mellerowicz et al., 2008; Nishikubo et al., 2007). Later studies reported no evidence of xyloglucan, and proposed pectins were responsible for generating tension in the G-layer rather than xyloglucan (Gorshkova et al., 2018; Guedes et al., 2017). Still others found xyloglucan in developing G-layers of Populus, with decreasing abundance at maturity (J. K. Kim & Daniel, 2019). How exactly G-fibers produce tensile force remains under investigation; however, a leading hypothesis involves the entrapment of matrix polysaccharides between laterally interacting cellulose microfibrils, producing longitudinal tensile stress (Chang et al., 2015; Gorshkova et al., 2018), consistent with *in vivo* visualizations showing expansion of the cellulose lattice during G-layer maturation (Clair et al., 2011).

G-fibers are well-characterized in the tension wood of angiosperm trees, where they form on the upper side of leaning tree trunks and generate sufficient tensile force to reorient growth towards the vertical: an example of negative gravitropism (Donaldson & Singh, 2016; Groover, 2016; Jourez et al., 2001; White & Robards, 1965). These fibers have also been identified in a variety of specialized organs, including the aerial roots of fig trees (Zimmermann et al., 1968), contractile roots of cycads (Tomlinson et al., 2014), fruit-bearing peduncles (Sivan et al., 2010), tendrils of climbing vines (Sousa-Baena et al., 2018), and are well-represented in the coiled stems of twiners (Bowling & Vaughn, 2009; Chery et al., 2022). To investigate the role of G-fibers in the twining growth habit, we sought to perturb their development. This has been accomplished in Poplar (*Populus sp*.) through application of exogenous Brassinosteroid phytohormones (BRs). In soil drench experiments, elevated BR levels promoted tension wood formation (and thus G-fibers), while BR inhibition impaired tension wood formation along with the plant’s gravitropic response (Du et al., 2020). In contrast, exogenous application of BRs via lanolin paste delayed G-fiber development and suppressed negative gravitropic bending, whereas BR inhibition markedly enhanced the stem’s gravitropic response (Gao et al., 2019).

The effect of BR application via lanolin paste on common bean was pronounced, disrupting the three major hallmarks of twining: internode elongation, circumnutation, and G-fiber deposition. BR treatment appeared to delay G-fiber maturation (consistent with Gao et al., 2019). Differential gene expression analysis highlighted a role for XTH (Xyloglucan endotransglucosylase/hydrolase) genes – which modify the hemicellulose, xyloglucan, in the cell wall, prompting us to test the controversial hypothesis that xyloglucan is present in the G-layer. Our findings support the idea that cell wall-remodeling XTH enzymes in both the primary wall and the G-layer contribute to the differentiation between the short, sturdy internodes typical during the shrubby growth phase and the elongated internodes characteristic of the twining phase in common bean.

## Materials and Methods

### Plant cultivation

This study utilized the Recombinant inbred line (RIL) L88-57 (Frahm et al., 2004), provided by Jonathan Lynch of The Pennsylvania State University. Seeds were primed in a 20% w/v solution of polyethylene glycol 4000 (Sigma Aldrich, Burlington, MA, USA) for 24 hours in the dark, rinsed thoroughly, and then placed on filter paper saturated in deionized water in Petri dishes for 48 hours under indirect light. Once germinated, seeds were transplanted into 10 cm plastic pots filled with Lambert LM-111 All Purpose Mix soil (Griffin Greenhouse Supplies, Tewksbury, MA, USA). A 500 ppm aqueous solution of Jacks 20-20-20 fertilizer (JR Peters, Allentown, PA, USA) was applied as needed. Plants were grown under 130 µmol m^−2^ s^−1^ PAR in Environmental Growth Chamber (EGC) Model GR48, with a 30-minute ramp up period, 12 hours of light, and a 30-minute ramp down to 12 hours of darkness, maintaining a relative humidity of 50%. Smooth, round plastic stakes (RooTrimmer, Nantong, China)—73 cm tall, 0.635 cm wide—were provided for plants to climb.

### Brassinosteroid treatment

One mg of the brassinosteroid 24-epibrassinolide (BL) (Goldbio, St Louis, MO) was dissolved in 1 ml of 95% v/v ethanol and thoroughly mixed into 100 g of melted lanolin (Sigma Aldrich). The same procedure was applied to the brassinosteroid-inhibitor brassinazole (BZ) (Sigma Aldrich). For the mock treatment, 1 ml of ethanol without hormones was added to the lanolin. These concentrations followed the method described in Gao et al., (2019). The hormone-lanolin paste was applied using a clean toothpick to the fourth internode of bean plants with five total internodes. The fourth internode was selected for treatment because it is typically the first to elongate and circumnutate, marking the transition from shrub to vine growth habit (Onyenedum et al., 2025).

Measurements of stem segments (hypocotyl, epicotyl, and internodes with mature trifoliate leaves) were taken before hormone treatment, three days after treatment, and again 18 days after treatment. There were between 13-21 biological replicates per each treatment and timepoint, for a total of 121 samples. This 18-day period corresponded to the mature, twined phenotype with ten trifoliate leaf-bearing internodes in the mock and BZ-treated plants. Although preliminary experiments showed that some BL-treated plants took longer to reach the same maturity benchmarks, an 18-day timeframe was selected as the standard to ensure consistency across treatments.

### Timelapse video

An Apple iPhone 12 (Cupertino, CA, USA) was used in conjunction with the Lapse It application (Interactive Universe Creative Softwares Eireli, http://www.lapseit.com) to capture one frame every 15 minutes. Frames taken in the dark were removed from the final timelapse.

### G-fiber measurement

To characterize the G-fibers, plants were harvested and stored in 70% ethanol three and 18 days after lanolin application. The fourth internodes were hand-sectioned using ASTRA01 razor blades (Gillette India, Rajasthan, India) and stained with 2% w/v toluidine blue (Sigma Aldrich) in water. Stained sections were viewed with bright-field optics using an Olympus BH2 microscope (Tokyo, Japan) and 10× /0.25 numerical aperture (NA), 20× /0.4 NA, or 40× /0.7 NA objective lenses. Images were taken using an AmScope MU 1000 digital camera (Irvine, CA, USA) mounted on the microscope. G-fibers were only present in plants 18 days after lanolin application.

G-layer thickness was measured from photographs captured with a 40× /0.7 NA objective lens. Images were processed with FIJI (Schindelin et al., 2012), and the line tool was used to measure 25 G-layers per pericyclic bundle. Each cross-section had between 12 and 14 bundles, with six biological replicates per treatment.

In order to measure G-fiber lengths, whole fourth internodes were macerated for three days in 70% nitric acid (Sigma Aldrich). The maceration solution was diluted with water, poured through a 40 µm cell strainer (Biologix, Shandong, China), and stained with 2% w/v toluidine blue. For each treatment, 25 G-fibers from six biological replicates were measured using AmScope software.

### RNA sequencing

Total RNA was isolated from treated fourth internodes at both timepoints using Qiagen’s RNeasy plant mini kit (Qiagen, Germantown, MD, USA) following the manufacturer’s instructions. There were ten biological replicates per treatment and timepoint (60 total samples). The RNA samples were then sent to Novogene Corporation Inc. (Sacramento, CA, USA) for library preparation, quality control, sequencing, and alignment. Novogene provided BAM files of aligned reads, gene-level count data, and quality control reports. For a detailed account of their methods see Supplementary File 1.

### Bioinformatics and visualization

RNA sequencing analysis was performed in R version 4.4.3 (R Core Team, 2025). BAM files received from Novogene were processed with the featureCounts function (Rsubread) to generate gene counts, which were filtered to retain genes with at least ten counts per million in at least one treatment group. Differential gene expression was assessed using DESeq2, with genes considered significantly differentially expressed if the log2 fold change was >1 and the adjusted p-value was <0.05. For the full dataset. Venn diagrams were generated using the VennDiagram package in R to illustrate the overlap of differentially expressed genes between treatments and timepoints. Gene Ontology (GO) terms were extracted from the *Phaseolus vulgaris* reference genome (v2.1, Phytozome) annotation file and mapped to descriptions using the GO.db package in R (Bioconductor). GO enrichment analysis was performed using a custom over-representation test with Fisher’s exact test, comparing the set of differentially expressed genes to the background set of expressed genes. Adjusted p-values were calculated using the Benjamini–Hochberg method. The top enriched GO terms were visualized as horizontal bar plots in ggplot2, ordered by statistical significance, with the x-axis representing the –log₁₀-transformed adjusted p-values. XTH genes were subsetted based on protein descriptions in the *Phaseolus vulgaris* reference genome. Variance stabilizing transformation (VST) was applied using DESeq2’s vst function, and counts were aggregated by treatment group using trimmed means. Volcano plots were generated to visualize the log2 fold changes versus the -log10 p-values of differentially expressed genes for each comparison and timepoint. Heatmaps were generated using the pheatmap package. For significantly DEGs, locusName (numeric gene ID from Phytozome) was matched to the homologous *Arabidopsis thaliana* protein name and symbol.

### Immunolocalization of xyloglucan

A blocking solution was prepared by dissolving 0.5 g of Carnation powdered milk (Vevey, Switzerland) in 10 ml of 1X Phosphate-Buffered Saline (PBS) in a 15 ml Falcon tube. Equal amounts of blocking solution were added to each 1.7 mL Eppendorf tube, and hand-sectioned stem cross-sections were incubated for 40 minutes. The samples were then washed three times with 1 ml 1X PBS for 5 minutes per wash.

Samples were incubated in a 1:100 dilution of Anti-Xyloglucan primary antibody (LM15, Megazyme) in 1X PBS for 90 minutes, followed by three washes with 1 ml PBS for 5 minutes each to remove nonspecific binding. Lastly, all samples were incubated in a 1:1000 dilution of Alexa 488 Goat anti-Rat lgG Fab fragment secondary antibody for 90 minutes, then washed three times with 1X PBS. Negative control samples were incubated in only secondary antibodies to observe non-specific fluorescence. Samples were mounted on slides using Citifluor Antifadent Mountant Solution (Hatfield, Pennsylvania, USA). Images were captured using a Leica Stellaris 5 Confocal Microscope (Leica Microsystems, Wetzlar, Germany) equipped with a 100X/1.4 NA oil immersion objective with a 488 nm excitation laser with a 525/50 nm emission filter for detection.

#### Quantification of LM15 signal

The GNU Image Manipulation Program (GIMP) was used to create masks defining the borders of the G-layer by manually tracing the cells in greyscale images. These masks were overlaid onto the original confocal images to delineate the G-layer boundaries.

The CellProfiler Speckle Count pipeline (available at Cellprofiler.org/examples) was utilized to distinguish smaller xyloglucan fluorescence signals from the larger gelatinous layer. This pipeline enabled quantification of the percentage of the total annotated area occupied by xyloglucan epitopes.

### Phylogenetic tree of XTH genes

A list of XTH genes was generated based on the *Phaseolus vulgaris* v2.1 annotation file from Phytozome (https://data.jgi.doe.gov/). These genes were BLASTed on NCBI (https://www.ncbi.nlm.nih.gov/) using BLASTN to obtain CDS Fasta files. The Fasta sequences were aligned with the MUSCLE alignment tool in AliView (Larsson, 2014). The maximum likelihood phylogenetic tree was created using RAxML-HPC2 (Stamatakis, 2014) (with 1000 bootstrap iterations via CIPRES (Miller et al., 2010) (https://www.phylo.org/), and visualized in FigTree v1.4.4 (Rambaut & Drummond, 2012) (https://tree.bio.ed.ac.uk/software/figtree/).

### In situ hybridization

Fourth internodes from plants 18 days after producing five sets of trifoliate leaves (equivalent to the 18 days after lanolin treatment, but without being treated) were fixed in FAA (3.7% formaldehyde, 5% acetic acid, 50% ethanol) for two hours, dehydrated in an ethanol series (70%, 85%, 95% ethanol for 1 hour each, 100% ethanol three times at one hour each), processed into toluene, embedded in Leica Surgipath® Paraplast Plus® Paraffin, and sectioned with a HM 315 microtome (Swerdlick Medical Systems, Simi Valley, CA) to a thickness of 10 µm. Paraffin sections were placed on Probe On Plus microscope slides (Fisher Scientific, Pittsburgh, PA).

Probes for in situ hybridization were created by extracting RNA using an RNeasy plant mini kit (Qiagen, Germantown, MD, USA) following the manufacturer’s instructions. RNA was treated with RNA Clean & Concentrator with DNase I (Zymo, Irvine, CA), and reverse transcribed using LunaScript RT SuperMix (New England BioLabs, Ipswich, MA). Gene specific fragments were generated by PCR using Phusion High Fidelity DNA polymerase (New England BioLabs) for XTH5 and HDzip (as a positive control gene to test if the hybridization worked). Primer sequences were as follows: XTH5—Forward primer: cacacagggcaagagatggt, Reverse primer: gtggcaactgccaagagaa, amplicon size 287 basepairs. PCR products were cleaned with a QIAQuick pcr purification kit (Quiagen) and cloned into chemically competent cells using the Zero Blunt TOPO PCR cloning kit (Thermo Fisher). Plasmids were extracted from transformed cells using the ZymoPURE plasmid miniprep kit (Zymo) and sent oK for sequencing to confirm the correct amplicon was present. The plasmid DNA was then used as the template for another PCR amplification using gene specific primers with T7 promoter sequences on the reverse primer were used to generate antisense probes. Fragments were cleaned with the Qiagen PCR cleanup kit and resuspended in 10uL of water. RNA probes containing Digoxigenin were generated according to the manufacturer’s (Sigma-Aldrich) protocol. In situ hybridization was performed as previously described (Ambrose et al., 2000; Ferrándiz et al., 2000) on three biological replicates. Images were captured using brightfield microscopy an Olympus BH-2 Microscope (Olympus Optical, Tokyo, Japan) at 20× and 40× objective magnifications.

### Statistics

An analysis of variance (ANOVA) was performed using R version 4.4.3 (R Core Team, 2025) to test for differences between the groups in G-fiber length and width, internode elongation, and speckle density of LM15 detection. Following significant results, paired t-tests were conducted for pairwise comparisons, with p-values adjusted using the Bonferroni method.

## Results

### BL treatment induces rapid localized elongation

Topically applied brassinolide (BL) in lanolin produced a notable elongation phenotype on the treated fourth internode (Fig. 1). This effect emerged within three days of the treatment (Fig. 1f) and was statistically significant for the BL-treated plants compared to the mock and BZ-treated plants, which were statistically indistinguishable (Fig. 1h). The lanolin was left on for the duration of the experiment, however a side experiment revealed the same elongation phenotype occurred even when the lanolin was removed after 24 hours (data not shown), suggesting the effect – at least on elongation – is likely the result of short-term signaling cascades, rather than prolonged exposure. The elongation that occurred on the fourth internode of BL-treated stems was never matched by the mock and BZ-treated plants; in fact, this elongated phenotype was even more pronounced at maturity (Fig. 1i-l). The elongation effect was only significant for the treated fourth internode, although insignificant elongation was observed for the fifth internode as well in BL-treated plants.

**Fig. 1:**
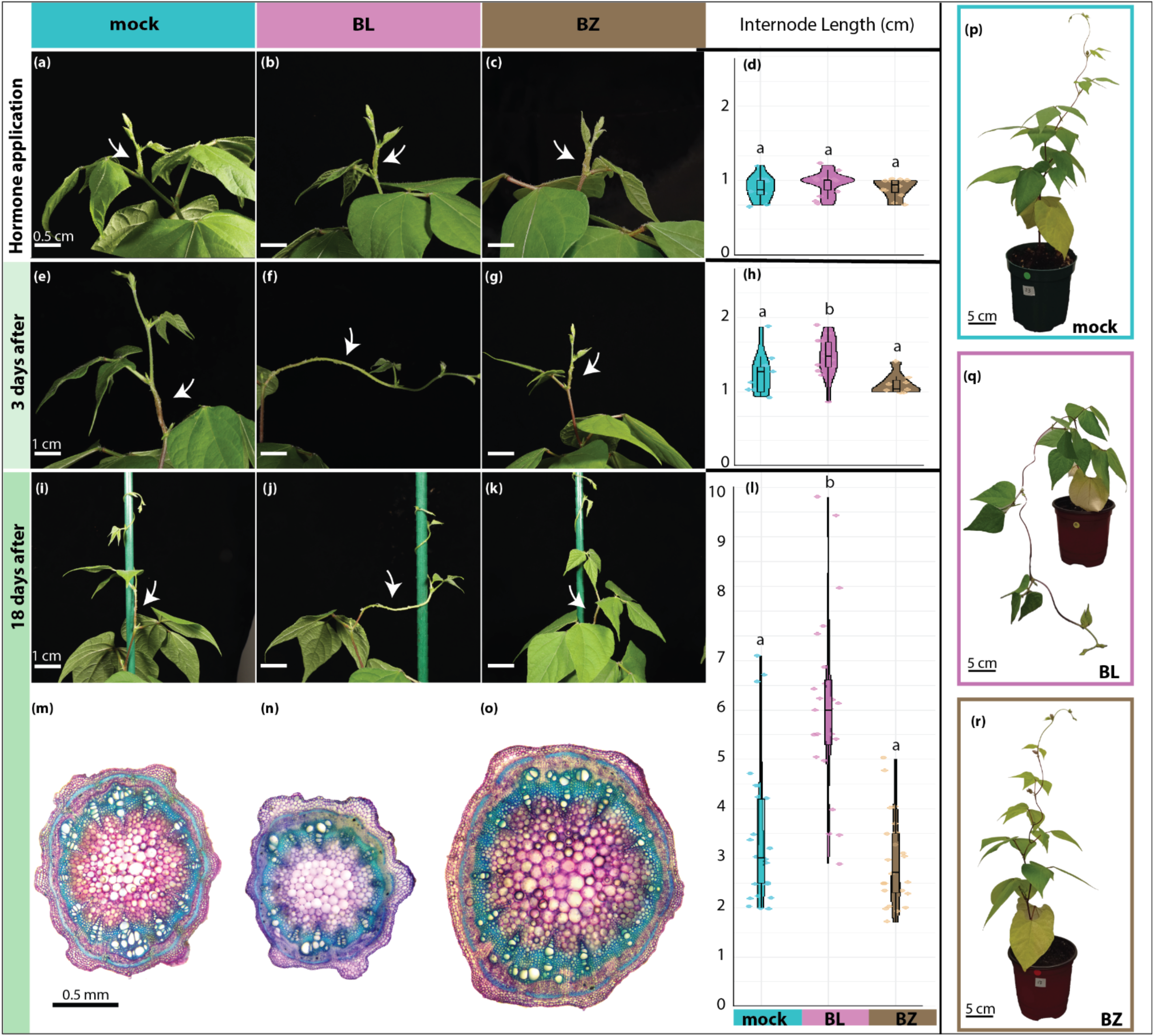
Effects of exogenous brassinosteroid manipulation on bean stem morphology and anatomy. Representative fourth Internode at the time of lanolin application for mock treatment (a), BL (24 epi-brassinolide, a brassinosteroid) treatment (b), and BZ (brassinazole, a brassinosteroid inhibitor) treatment (c). Plot showing internode lengths at the time of lanolin application (d). Representative fourth internode three days after lanolin application for mock treatment (e), BL treatment (f), and BZ treatment (g). plot showing internode length three days after lanolin application (h). Representative fourth internode 18 days after lanolin applications for mock treatment (i), BL treatment (j), and BZ treatment (k). Plot showing internode lengths 18 days after treatment (l). Representative cross-section of the treated fourth internode for mock treatment (m), BL treatment (n), and BZ treatment (o). Whole-plant phenotype 18 days after treatment with stake removed for mock treatment (p), BL treatment (q), and BZ treatment (r). For plots (d, h, l) statistical analysis was performed using the Kruskal-Wallis test to compare differences among groups at each timepoint, followed by pairwise comparisons with Bonferroni-adjusted p-values. Groups with p < 0.05 were considered statistically significant and denoted with letters. n ≥13 biological replicates per treatment and time point.

### BL treatment impedes circumnutation, twining, and mature stem stability

The BL-treated internodes were narrower compared to mock and BZ-treated internodes (Fig. 1 m–o). Notable asymmetrical cell expansion of the cortex was also observed in BL-treated cross-sections (Fig. 1n). The result of localized elongation on the fourth internode produced whole-stem instability for mature BL-treated plants: while mock and BZ-treated stems were able to maintain an erect posture at maturity after their stake was removed, the BL-treated plants flopped over when the stake was removed (Fig. 1q). This elongation also impeded the BL-treated plants’ ability to circumnutate, find a support, and twine, at least from the treated internode. While mock and BZ-treated internodes elongated vertically (Fig. 1e, 1g), BL-treated internodes exhibited a loss of structural stability, bending to one side as they elongated, resulting in a horizontally oriented fourth internode (Fig. 3f).

Timelapse videos of the plants showed that the BL-treated internodes produced erratic circumnutational movements (Video 1). Typical smooth circumnutation resumed in upper internodes not treated with BL, although this either delayed twining or led to plants being unable to successfully locate a stake entirely.

### BR treatment results in the development of longer, thinner G-fibers

Mock, BL, and BZ-treatment on the fourth internode produced three morphologically and statistically distinct groups of G-fibers, which mirrored the effects of brassinosteroid manipulation on the internodes as a whole (Fig. 2). To measure G-fiber thickness, six internodes per treatment were cross-sectioned, and 25 cells from each pericyclic bundle (12-14) were measured (n ≈ 1800 cells per treatment) to account for potentially asymmetric distribution of G-fiber thickness. The BL-treated internodes produced G-fibers that were significantly thinner than the other groups (Fig. 2b). By contrast, the BZ-treated internodes produced G-fibers that were thicker than the other groups (Fig. 2c). The mock-treated internodes were intermediate in terms of thickness (Fig. 2a), and statistically distinct from both the thin BL-treated G-fibers and thick BZ-treated G-fibers (Fig. 2d). Whole internodes were macerated in nitric acid to isolate G-fibers for longitudinal measurements (Fig. 2e). Three biological replicates were made, with 25 G-fibers measured from each (n=75 cells per treatment). Again, the three treatments fell into three statistically significantly different groups: the BL-treated group was the longest, averaging 6.27 mm; the BZ-treated group was the shortest, averaging 3.22 mm; the mock-treated group averaged an intermediate 4.14 mm (Fig. 2f).

**Fig. 2:**
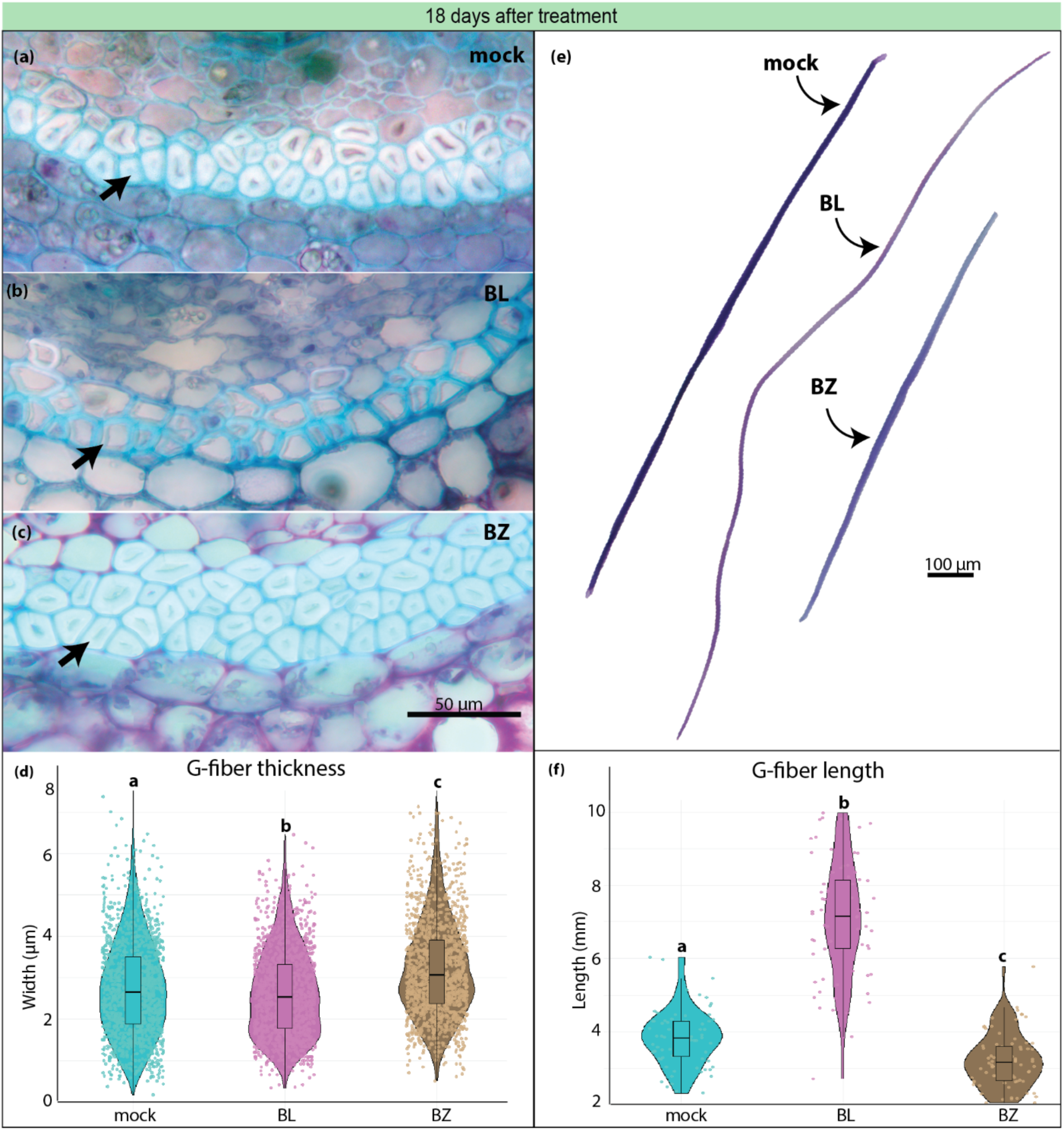
Differences in G-fiber morphology between treatment groups 18 days after lanolin application. Representative cross-sections of plants that received mock (a), BL (b), and BZ (c) treatment. Arrows indicate G-fibers. Plot showing the G-fibers widths by treatment from six biological replicates (d). Representative G-fibers isolated from tissue maceration of the reated fourth internode (e). Plot showing the G-fiber lengths by treatment from three biological replicates (f)). Letters denote statistically significant groups, p > 0.5.

### XTH genes are diMerentially expressed between treatment groups

To examine the genetic basis of the above phenotypes, we measured gene expression across the genome in mock-, BL-, and BZ-treated fourth internode, three days and 18 days after treatment. Differential gene expression (DGE) was found between all treatment groups (Fig. 3). After filtering out low/inconsistently expressed genes, the dataset included 12,587 genes. In general, there were more DEGs three days after treatment (when the differential phenotype was beginning to emerge), compared to 18 days after treatment (when the plants were mature). The greatest difference was between the BL and BZ treatments, with the highest level of DGE occurring three days after treatment; 476 (3.8%) genes were upregulated and 570 (4.5%) genes downregulated in BL compared to BZ. The imperceptible phenotypic difference upon BZ treatment compared to mock treatment after three days was reflected in the differential expression of only 30 genes in total; however, the number of DEGs for this comparison increased later on, 18 days after treatment, to 155 genes.

**Fig. 3:**
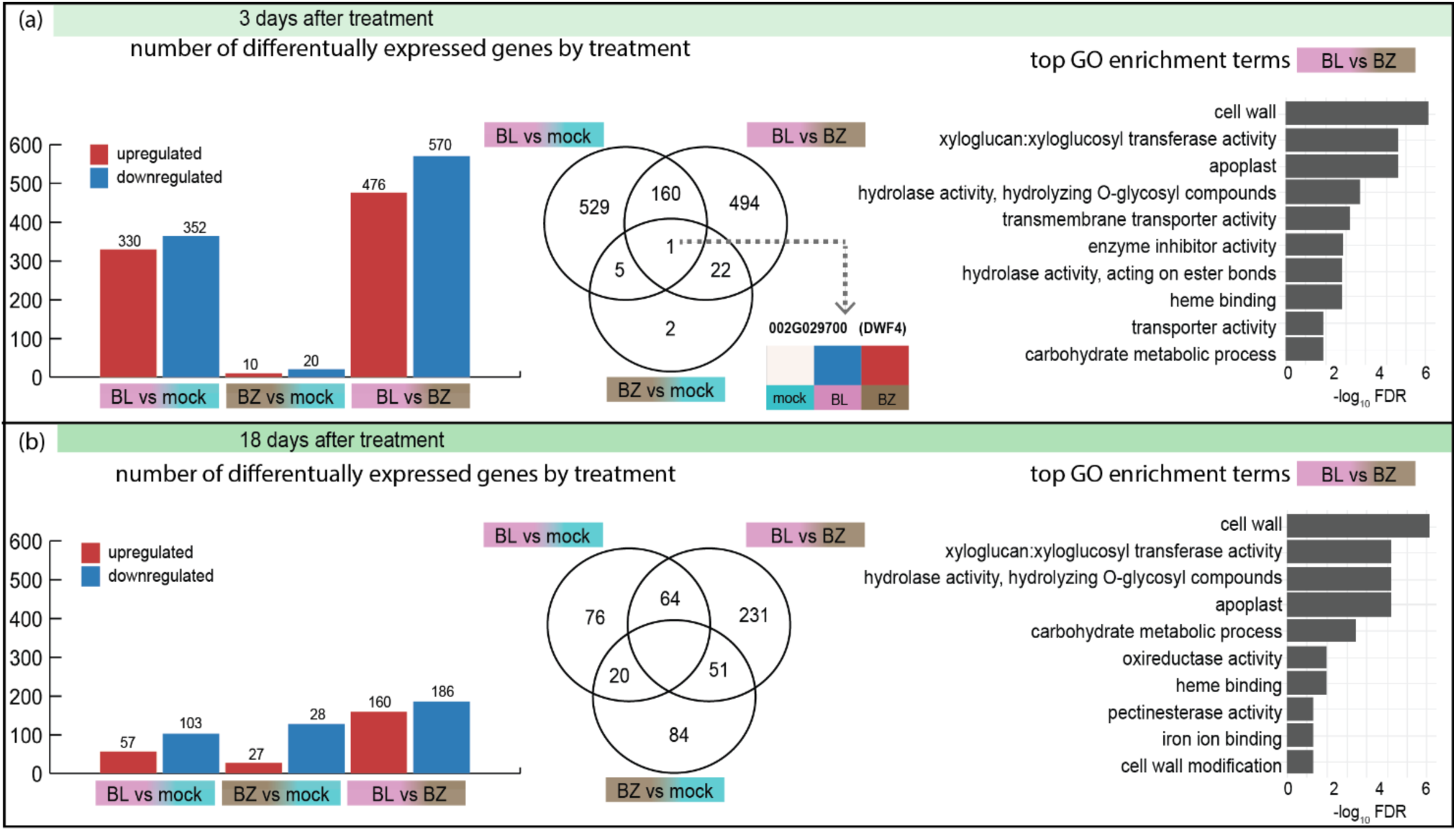
Differential Gene Expression. Bar graph showing number of differentially expressed genes by treatment, venn diagram showing the number of differentially expressed genes by each treatment comparison, and top GO enrichment terms for comparison between brassinolide (BL) and brassinazole (BZ) treatment, three days after treatment (a), and 18 days after treatment (b), indicating the major functions of the top differentially expressed genes.

As shown in the Venn diagrams for each timepoint in Fig. 3, only one gene was differentially expressed across all treatment comparisons, and only at the three days after treatment timepoint. This gene, DWF4 (Phytozome locusName: 002G029700), is a key component of BR biosynthesis. No brassinosteroid-related genes were differentially expressed 18 days after lanolin application, indicating that the BR perturbation was established early. Expression of DWF4 was downregulated in BL-treated plants and upregulated in BZ-treated plants relative to mock treatment (Fig. 3a).

Since the greatest differential gene expression occurred between the BL and BZ treatments—representing brassinosteroid application and inhibition, respectively—this comparison was used to identify the most significantly enriched GO terms. Across both time points, the top four enriched terms were cell wall (GO:0005618), xyloglucan:xyloglucosyl transferase activity (GO:0016762), apoplast (GO:0048046), and hydrolase activity, hydrolyzing O-glycosyl compounds (GO:0004553) (Fig. 3). Notably, xyloglucan-modifying XTH genes were associated with all four GO terms and comprised 100% of the genes annotated with both the xyloglucan:xyloglucosyl transferase activity and apoplast terms. Of the 112 differentially expressed genes annotated with the cell wall term in the reference genome (v2.1, Phytozome), 76 encode pectin-modifying enzymes, 33 are involved in xyloglucan modification, and 3 encode serine proteases. The hydrolase activity, hydrolyzing O-glycosyl compounds term included a mix of glycosyl hydrolases and glucosidases.

Due to incomplete genome annotation, other gene families known to contribute to G-fiber development, such as the fasciclin-like arabinogalactan proteins (FLAs) (Lafarguette et al., 2004), were not assigned GO terms. However, several FLA genes (FLA2, FLA7, FLA9, and FLA16) were significantly upregulated in BL-treated internodes compared to BZ-treated internodes, and FLA2 and FLA9 were significantly upregulated in BL-treated internodes compared to mock treatment.

Among the significant DEGs potentially impacting the phenotypic changes via cell wall remodeling, XTH (xyloglucan endotransglycosylase/hydrolase) genes emerged as candidates of particular interest. XTH genes were among the most highly differentially expressed in initial exploratory datasets and were annotated with all four of the top GO enrichment terms for DEGs between the BL and BZ treatments at both time points (Fig. 3). XTH genes were significantly differentially expressed in BL-treated internodes compared to both mock and BZ-treated internodes at both three and 18 days after treatment (Fig. 4a-f), potentially corresponding to both primary cell wall and G-layer remodeling, respectively. Most XTH genes in mature BL stems were relatively downregulated, possibly indicating ongoing deposition.

**Fig. 4:**
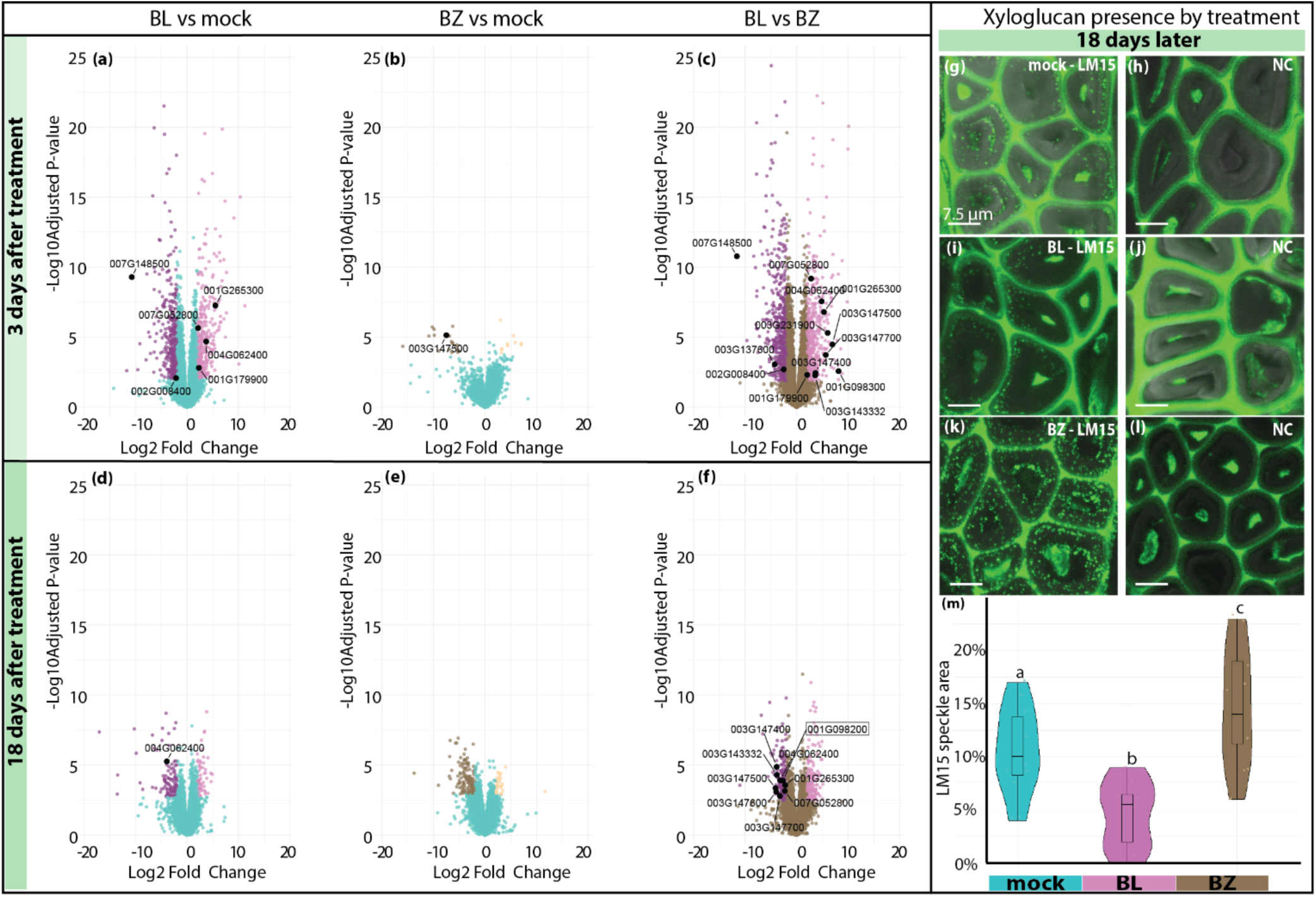
Differential Gene Expression (DGE) between treatment groups and xyloglucan presence in the G-layer. Volcano plots of DGE between treatments (panels a-f). Each point on the volcano plots represents a gene. Genes that do not significantly differ in expression between the two treatments are in the same color as the baseline treatment (turquoise for mock, brown for BZ). Differentially expressed genes (log2 fold change > 1 and adjusted p-value < 0.05) appear in the colors of the comparison treatment (pinkish hues for BL, brownish hues for BZ). Upregulated genes appear on the right and downregulated genes appear on the left. Significant XTH genes are highlighted in black and labeled with their Phytozome locusNames. Specific comparisons are as follows: (a) BL vs. mock, (b) BZ vs. mock, (c) BL vs. BZ at 3 days after treatment; (d) BL vs. mock, (e) BZ vs. mock, (f) BL vs. BZ at 18 days after treatment. Presence of xyloglucan in the G-layer (panels g–m). Confocal microscopy images of cross-sections from treated fourth internodes stained with the monoclonal antibody LM15. Xyloglucan appears as speckles within the G-layer. Speckles are visible in the mock-treated (g), BL-treated (i), and BZ-treated (k) internodes. No speckles are observed in the corresponding negative controls—mock NC (h), BL NC (j), and BZ NC (l). Panel (m) shows quantification of LM15 speckle density across treatments. Statistical analysis was performed using ANOVA followed by pairwise t-tests with Bonferroni-adjusted p-values. Groups with p < 0.05 were considered statistically significant and are denoted by different letters.

### Xyloglucan presence confirmed in the G-layer

The potential for XTH genes to contribute to the observed phenotype depends on whether these genes act on the G-layer or are limited to the primary cell wall. To assess their possible involvement in G-layer deposition, immunostaining with a xyloglucan-detecting monoclonal antibody, LM15, was performed to test for the presence of xyloglucan in the G-layer (Fig. 4g-l). Not only was xyloglucan clearly present under all treatments, but its presence fell into three distinct groups: moderate presence in mock-treated internodes (Fig. 4g), reduced presence in BL-treated internodes (Fig. 4i), and elevated presence in BZ-treated internodes (Fig 4k). Image analysis was employed to quantify these differences, and the results showed three statistically significantly different groups based on the area of LM15 speckles relative to the area of the G-layer (Fig. 4m).

### Expression patterns of Phaseolus vulgaris XTH genes vary by time and treatment

Once it was determined that XTH genes are differentially expressed based on BR-treatment and confirmed that xyloglucan is present in the G-layers of common bean, the next task was identifying which XTH genes might be responsible for the observed G-fiber differences. There are 34 genes in the *Phaseolus vulgaris* genome annotated as XTH, many of which showed differential expression. To better understand the relationship of these genes to each other and their expression patterns through development, a phylogeny of XTH genes was generated and paired with a heatmap of their corresponding expression by treatment and time point (Fig. 5). Of the 33 XTH genes, 13 were absent from the RNA extracted from the treated fourth internodes. These genes appear as grey cells in the heatmap and are likely expressed in other tissue types or at another developmental time point. The remaining XTH genes were typically primarily upregulated either three days after treatment or 18 days after treatment. This likely corresponds to XTH genes acting primarily on extending the primary cell wall to establish the elongation phenotype (upregulated after three days) and those which might play a continuing role in G-fiber development (upregulated after 18 days). There were a few exceptions, such as XTH15 (001G265300) and XTH5 (001G098300), which were upregulated in BL but downregulated in mock and BZ three days after treatment, and had the reverse pattern 18 days after treatment (Fig. 5).

**Fig. 5:**
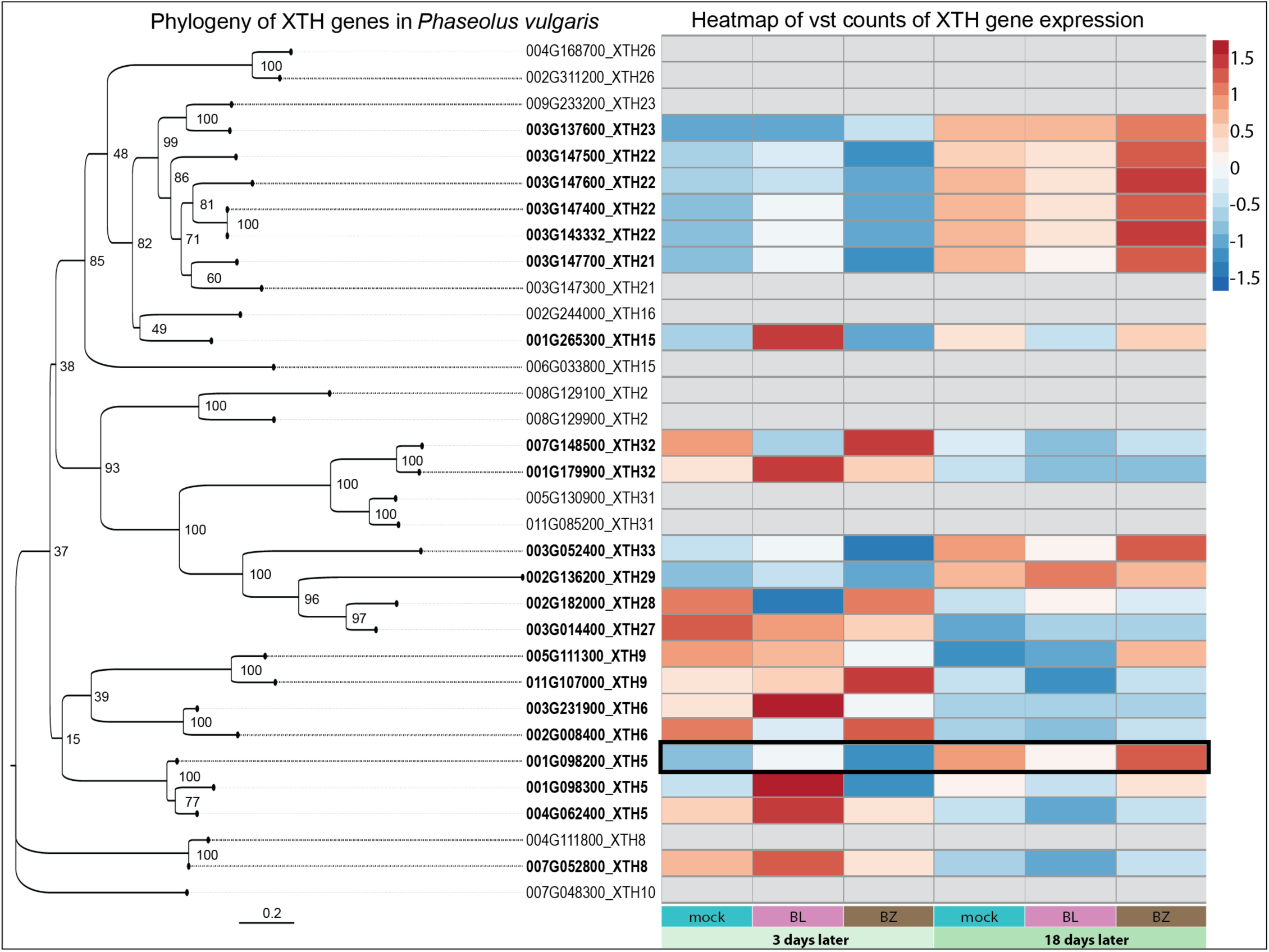
Phylogeny and Heatmap of Phaseolus vulgaris XTH genes. Unrooted phylogeny based on nucleic acid sequences of XTH genes in the Phaseolus vulgaris reference genome, constructed using maximum likelihood (1000 bootstraps) and paired with a heatmap depicting differential expression in the treated fourth internode over time. Gene expression values were derived from vst-transformed counts and aggregated by treatment group using 10% trimmed means. The color scale represents relative gene expression, with red indicating upregulation and blue indicating downregulation, based on Z-scores calculated by scaling each gene’s expression values across treatment groups. Gene IDs are annotated with their corresponding products, and ordering reflects the phylogenetic relationships among genes.

In selecting a candidate gene to test whether XTH activity was occurring in the G-layer, the criteria were: 1) The gene was upregulated during G-fiber development (18 days after treatment only) to avoid targeting a XTH gene related to primary cell walls; 2) The sequence pairwise differences were enough to avoid challenges with primer design; and 3) The gene was significantly differentially expressed across BR-treatments. While XTH21– XTH23 showed promise individually, their phylogenetic relatedness, pairwise sequence similarity and expression patterns hinted at potential genetic redundancy (Fig. 5). XTH33 displayed the right expression pattern, however the difference in expression was not statistically significant between groups. This left XTH5 (001G098200), which was upregulated 18 days after treatment, with a distinct sequence from phylogenetic sisters, and was significantly differentially expressed in BL versus BZ treated internode (Fig. 4f). XTH5 expression patterns and location in the phylogeny can be seen in Fig. 5, where the expression data are highlighted by a black box.

### XTH5 expression occurs within mature G-fibers

XTH5 (001G098200) was selected as a candidate gene for assessing spatial XTH expression in tissues producing G-fibers. In situ hybridization with an antisense RNA probe revealed strong, localized signal in G-fibers with well-developed G-layers in mock plants (Fig. 6a, b). A sense probe was used as a control and produced no detectable signal in the G-layer (Fig. 6c, d), indicating that the observed signal represents specific expression of XTH5.

**Fig. 6:**
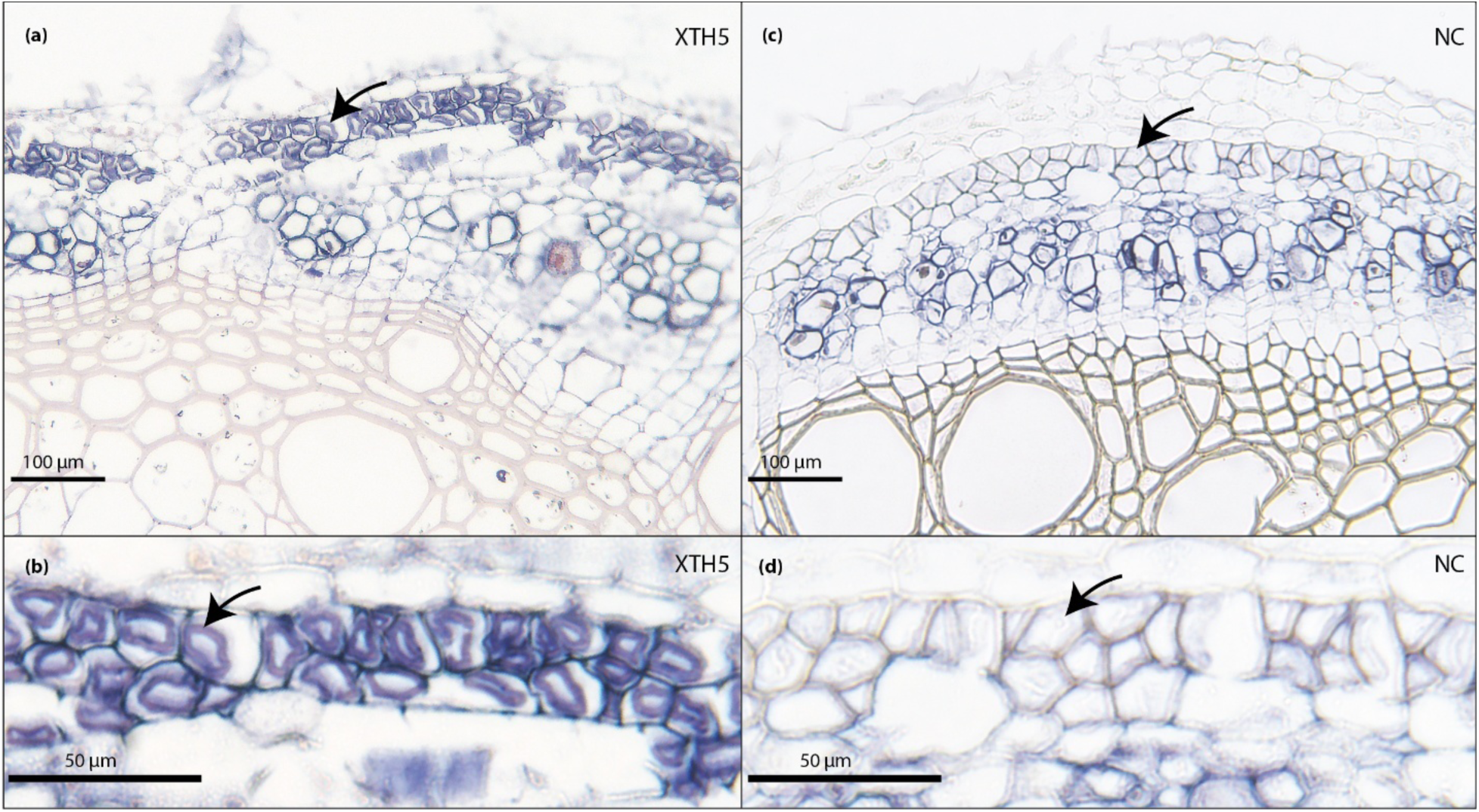
In situ hybridization of XTH5 transcripts in the G-fibers. Detection of XTH5 mRNA using a DIG-labeled antisense probe is visible as a purple precipitate localized to the G-fibers, visualized under brightfield microscopy at 20× (a) and 40× (b). No signal is observed in the negative control (NC) sections hybridized with a DIG-labeled sense probe, shown at 20× (c) and 40× (d). Black arrows point to G-layer.

## Discussion

Our study extends G-fiber research into a vine species, common bean, that forms G-fibers in association with its twining habit. Notably, BR treatment in common bean impacted all three characteristics of the twining phenotype (i.e., the shifts that mark the transition between shrub or vine growth habit): elongation (Fig. 1), exaggerated circumnutation (Video 1), and G-fiber deposition (Fig. 2). Previous experimental results involving BR manipulation on G-fibers in trees varied with dosage and application method. Du et al. reported that elevated BR levels, introduced through soil drench or genetic modification, enhanced tension wood (and thus G-fiber) formation (Du et al., 2020). In contrast, Gao et al. found that lanolin-applied BRs promoted G-fiber elongation but delayed maturation (Gao et al., 2019). Our results align with the findings of Gao et al.(2019): BL-treated fibers were more elongated, but thinner, than controls (Fig. 2). We found that the effect was localized to the treated area: internodes and G-fibers above and below the treated region were not impacted. This is consistent with reports that native BRs act locally in plant organs and do not undergo long-distance transport (Symons & Reid, 2004; Vukašinović et al., 2021).

The rapid onset of internode elongation following BL treatment (Fig. 1) is consistent with the known function of brassinosteroids in promoting cell expansion and elongation, with cellular-level effects reported within minutes of application (Caesar et al., 2011). At the first RNA sequencing timepoint (three days after treatment, when elongation begins) *DWF4* was the only gene significantly differentially expressed across all three treatment comparisons (Fig. 3a). *DWF4* encodes a cytochrome P450 enzyme responsible for key 22α-hydroxylation steps in BR biosynthesis and is a rate-limiting component of the pathway (Choe et al., 1998; Fu et al., 2023; Zebosi et al., 2024). Because *DWF4* transcription is negatively regulated by BR levels as part of a well-characterized feedback loop, its expression serves as an indicator of BR signaling activity (Yoshimitsu et al., 2011). BR-deficient or insensitive plants often show reduced growth due to impaired cell expansion. For example, *dwarf4* (*cyp90*) *Arabidopsis* mutants exhibit shortened hypocotyls, male sterility, and stunted growth, all of which can be rescued by exogenous BL application (Choe et al., 1998; Szekeres et al., 1996). In our study, *DWF4* was strongly downregulated in BL-treated plants, consistent with elevated BR levels, and significantly upregulated in BZ-treated plants, indicating a compensatory response to reduced endogenous BR. Although we did not sample RNA immediately after treatment to capture rapid changes in gene expression, the differential expression of *DWF4* after three days confirms successful manipulation of endogenous BR levels.

Differentially expressed genes associated with the top GO terms (Fig. 3) included cell wall-modifying genes such as BGALs (Beta-Galactosidases), XTHs, and PMEs/PMEIs (Pectin Methylesterase/Pectin Methylesterase Inhibitors) consistent with transcriptional programs observed during G-fiber development in other species. BGALs and XTHs are upregulated in constitutive G-fiber plants like ramie (*Boehmeria nivea*) and flax (*Linum usitatissimum*), as well as during tension wood formation in poplar (*Populus*) (Sousa-Baena & Onyenedum, 2022). This is consistent with upregulation of these genes in BZ-treated internodes with seemingly more mature G-fibers (Fig. 4).

Another gene family typically upregulated during G-fiber development in other species are the FLAs (Fasciclin-Like Arabinogalactan proteins) (Takeuchi et al., 2024; Wang et al., 2017). Several FLAs showed increased expression (FLA2 and FLA9 were significantly upregulated in BL-treated internodes compared to mock treatment), but none were differentially expressed at the time of G-fiber deposition (18 days).

### Xyloglucan is Present in the G-layer of Common Bean

The prominence of XTH genes among top differentially expressed genes between BR treatment groups prompted us to investigate whether their substrate, xyloglucan, is present in the G-layer of common bean. While xyloglucan is the most abundant hemicellulose in eudicot primary cell walls (Scheller & Ulvskov, 2010), its remodeling is generally thought to end with the cessation of cell expansion and the onset of secondary wall deposition, where xyloglucan is typically absent. The expression of XTH genes in mature internodes suggests continued re-arrangement of xyloglucan crosslinks, possibly within developing G-layers of pericyclic fibers, contributing to the observed differential phenotypes.

The presence of xyloglucan in the G-layer is supported by earlier studies on poplar tension wood, which found xyloglucan to be the most abundant non-cellulosic component of G-layers—comprising up to 15% of their molecular composition (Nishikubo et al., 2007). Xyloglucan has been theorized to be integral to the tensile properties of G-fibers and was referred to as the “molecular muscle of trees” (Mellerowicz et al., 2008). This hypothesis posits that xyloglucan could be trapped by adjacent cellulose microfibrils in the G-layer, resulting in longitudinal tensile stress as the cellulose lattice structure expands to accommodate encapsulated xyloglucan. Xyloglucan is also hypothesized to serve as a molecular tether between the tension-generating G-layer and the surrounding lignified secondary wall, facilitating stress transfer across these layers. Supporting this model, XTH genes are upregulated in developing tension wood compared to normal wood, and the associated XTH enzymes persist for years in mature G-fibers, dynamically adjusting xyloglucan–cellulose interactions between the G-layer and secondary cell wall (Nishikubo et al., 2007).

However, an investigation in 2017 using CCRC-M1 antibodies (the same as Nishikubo et al.) did not detect xyloglucan in the G-layers of poplar trees and posited rhamnogalacturonan pectin and arabinogalactan proteins as more likely candidates responsible for tension wood properties (Guedes et al., 2017). A later report *did* detect xyloglucan in the G-layers of poplar using LM15, but noted that its presence appeared to diminish as the G-fibers matured (J. K. Kim & Daniel, 2019). One explanation for these discrepancies is the hypothesis that xyloglucan migrates from the plasma membrane through the G-layer to its final location at the interface of the G-layer and secondary cell wall, supported by observations that xyloglucan is mainly detected at the external boundary of the G-layer (Baba et al., 2009). This would explain why xyloglucan is more readily detected in young G-fibers. Another explanation is that detection (rather than presence) is reduced in mature G-fibers, either as xyloglucan is modified by XTH activity making it unrecognizable to antibodies, or as cellulose aggregation entraps xyloglucan, making it inaccessible to antibodies (J. K. Kim & Daniel, 2019).

In our study, LM15 labeling was strong along the G-layer/secondary cell wall interface, particularly in fibers where the G-layer had become slightly detached during sectioning, consistent with (Baba et al., 2009). However, our imaging also indicates that xyloglucan is distributed throughout the entire G-layer, as suggested in (J. K. Kim & Daniel, 2019; Nishikubo et al., 2007), not merely restricted to the interface between the G-layer and secondary cell wall. Furthermore, the density of LM15 labeling varied significantly between treatment groups: it was lowest in the thin, elongated G-fibers of BL-treated internodes and highest in the thick, truncated G-fibers of BZ-treated internodes (Fig. 4). It remains unclear whether differences in G-fiber thickness reflect developmental timing: for instance, BL-treated G-fibers might require more time to reach full maturity, whereas BZ-treated fibers might thicken more rapidly due to their shorter length. Our results suggest that more developed G-fibers in BZ-treated internodes exhibit greater xyloglucan detectability via LM15.

### XTH Gene Expression Patterns Might Reflect Function

Patterns of XTH expression revealed that individual genes were primarily upregulated either at the early (three days) or late (18 days) timepoint, suggesting temporal specialization of different XTH isoforms. We hypothesize that early-expressed XTHs function in primary cell wall expansion and elongation (Van Sandt et al., 2007), whereas later-expressed XTHs might act on xyloglucan within the G-layer, contributing to its tensile properties. This idea of early-elongation XTH genes is supported by work on cotton fibers (*Gossypium*), where early XTH expression has been linked to fiber elongation (Ji, 2003). Manipulating BR levels in cotton produces comparable outcomes: BL promotes fiber elongation, while BZ inhibits it, accompanied by differential expression of genes including XTHs (Sun et al., 2005; Yang et al., 2014).

### In-situ Hybridization Indicates Xyloglucan Remodeling in G-fibers

To further assess the role of XTH, we employed in situ hybridization to determine spatial expression. Specifically, we aimed to test whether XTH genes in mature internodes were expressed within the G-fibers, and if so, whether it is localized to the interface between the G-layer and secondary cell wall or more widely distributed. An alternative hypothesis is that the late-expressed XTH genes are unrelated to G-fiber development and instead serve other functions in mature internodes. Notably, XTH gene family members have been implicated in a range of abiotic stress responses, e.g., flooding (Song et al., 2018), salinity and drought (Cho et al., 2006; Choi et al., 2011), freezing (Takahashi et al., 2021), and heavy metals (Xuan et al., 2016), raising the possibility that their expression at later stages is linked to something other than G-layer remodeling. XTH5 was selected as a candidate for in situ hybridization based on its significant upregulation at 18 days and its differential expression across treatments: lowest in BL, intermediate in mock, and highest in BZ (Fig. 5). In situ hybridization revealed XTH5 expression distributed throughout the G-fiber (Fig. 6), supporting the conclusion that XTH genes, like xyloglucan itself, are integral to G-fiber development.

This spatial pattern of xyloglucan throughout the G-layer alongside G-fiber specific XTH activity supports the idea that xyloglucan plays a broader structural role within the G-layer. In particular, it may contribute to the organization of cellulose microfibrils (Bou Daher et al., 2024). We propose that xyloglucan not only mediates the attachment of the G-layer to the lignified secondary wall but also reinforces the internal structure of the G-layer, thereby facilitating effective tension distribution across the fiber.

## Conclusion

Exogenous brassinosteroid treatment induced rapid internode elongation, disrupted circumnutation, and resulted in G-fibers that were significantly longer and thinner than those of mock or inhibitor treated plants. XTH genes were mostly upregulated early (three days after treatment) in the brassinosteroid treatment, corresponding to elongation of the treated internode compared to mock and inhibitor treatment. Later, (18 days after treatment) XTH upregulation was more associated with the mature G-fiber phenotype, prominent in inhibitor and mock treated plants. The presence of xyloglucan as a constituent of the G-layer of common bean was established, as well as the expression of a representative gene, XTH5, localized via in situ hybridization to be expressed within the G-fiber itself. Together, these findings support a model in which XTH-mediated remodeling of xyloglucan contributes to both early cell expansion and later G-fiber development, positioning XTH genes as key regulators of twining-associated traits in common bean.

## Supplemental Material

### Supplementary File 1

Methods as provided by Novogene

### 1. Experimental Procedure

#### 1.1 Sample quality control

RNA integrity was assessed using the Bioanalyzer 2100 system (Agilent Technologies, CA, USA).

#### 1.2 Library preparation for Transcriptome sequencing

Messenger RNA was purified from total RNA using poly-T oligo-attached magnetic beads. After fragmentation, the first strand cDNA was synthesized using random hexamer primers followed by the second strand cDNA synthesis. The library was ready after end repair, A-tailing, adapter ligation, size selection, amplification, and purification. The library was checked with Qubit and real-time PCR for quantification and bioanalyzer for size distribution detection. Strand specific library Messenger RNA was purified from total RNA using poly-T oligo-attached magnetic beads. After fragmentation, the first strand cDNA was synthesized using random hexamer primers. Then the second strand cDNA was synthesized using dUTP, instead of dTTP. The directional library was ready after end repair, A-tailing, adapter ligation, size selection, amplification, and purification. The library was checked with Qubit and real-time PCR for quantification and bioanalyzer for size distribution detection.

#### 1.3 Clustering and sequencing

After library quality control, different libraries were pooled based on the effective concentration and targeted data amount, then subjected to Illumina sequencing. The basic principle of sequencing is "Sequencing by Synthesis", where fluorescently labeled dNTPs, DNA polymerase, and adapter primers are added to the sequencing flow cell for amplification. As each sequencing cluster extends its complementary strand, the addition of each fluorescently labeled dNTP releases a corresponding fluorescence signal. The sequencer captures these fluorescence signals and converts them into sequencing peaks through computer software, thereby obtaining the sequence information of the target fragment.

### 2. Bioinformatics Analysis Pipeline

#### 2.1 Data quality control

Raw data (raw reads) of fastq format were firstly processed through fastp software. In this step, clean data (clean reads) were obtained by removing reads containing adapter, reads containing ploy-N and low quality reads from raw data. At the same time, Q20, Q30 and GC content the clean data were calculated. All the downstream analyses were based on the clean data with high quality.

#### 2.2 Reads mapping to the reference genome

Reference genome and gene model annotation files were downloaded from genome website directly. Index of the reference genome was built using Hisat2 v2.0.5 and paired-end clean reads were aligned to the reference genome using Hisat2 v2.0.5. We selected Hisat2 as the mapping tool for that Hisat2 can generate a database of splice junctions based on the gene model annotation file and thus a better mapping result than other non-splice mapping tools.

#### 2.3 Novel transcripts prediction

The mapped reads of each sample were assembled by StringTie (v1.3.3b) (Mihaela Pertea.et al. 2015) in a reference-based approach. StringTie uses a novel network flow algorithm as well as an optional de novo assembly step to assemble and quantitate fulllength transcripts representing multiple splice variants for each gene locus.

#### 2.4 Quantification of gene expression level

featureCounts v1.5.0-p3 was used to count the reads numbers mapped to each gene. And then FPKM of each gene was calculated based on the length of the gene and reads count mapped to this gene. FPKM, expected number of Fragments Per Kilobase of transcript sequence per Millions base pairs sequenced, considers the effect of sequencing depth and gene length for the reads count at the same time, and is currently the most commonly used method for estimating gene expression levels.

## Notes

### Competing Interest Statement

The authors have declared no competing interest.

### Funding

- National Science Foundation: 2401675

## References

Ambrose, B. A., Lerner, D. R., Ciceri, P., Padilla, C. M., Yanofsky, M. F., & Schmidt, R. J. (2000). Molecular and Genetic Analyses of the *Silky1* Gene Reveal Conservation in Floral Organ Specification between Eudicots and Monocots. Molecular Cell, 5(3), 569–579. 10.1016/S1097-2765(00)80450-5

Baba, K., Park, Y. W., Kaku, T., Kaida, R., Takeuchi, M., Yoshida, M., Hosoo, Y., Ojio, Y., Okuyama, T., Taniguchi, T., Ohmiya, Y., Kondo, T., Shani, Z., Shoseyov, O., Awano, T., Serada, S., Norioka, N., Norioka, S., & Hayashi, T. (2009). Xyloglucan for Generating Tensile Stress to Bend Tree Stem. Molecular Plant, 2(5), 893–903. 10.1093/mp/ssp054

Bou Daher, F., Serra, L., Carter, R., Jönsson, H., Robinson, S., Meyerowitz, E. M., & Gray, W. M. (2024). Xyloglucan deficiency leads to a reduction in turgor pressure and changes in cell wall properties, affecting early seedling establishment. Current Biology, 34(10), 2094–2106.e6. 10.1016/j.cub.2024.04.016

Bowling, A. J., & Vaughn, K. C. (2008). Immunocytochemical characterization of tension wood: Gelatinous fibers contain more than just cellulose. American Journal of Botany, 95(6), 655–663. 10.3732/ajb.2007368

Bowling, A. J., & Vaughn, K. C. (2009). Gelatinous fibers are widespread in coiling tendrils and twining vines. American Journal of Botany, 96(4), 719–727. 10.3732/ajb.0800373

Caesar, K., Elgass, K., Chen, Z., Huppenberger, P., Witthöft, J., Schleifenbaum, F., Blatt, M. R., Oecking, C., & Harter, K. (2011). A fast brassinolide-regulated response pathway in the plasma membrane of Arabidopsis thaliana. The Plant Journal, 66(3), 528–540. 10.1111/j.1365-313X.2011.04510.x

Chang, S.-S., Quignard, F., Alméras, T., & Clair, B. (2015). Mesoporosity changes from cambium to mature tension wood: A new step toward the understanding of maturation stress generation in trees. New Phytologist, 205(3), 1277–1287. 10.1111/nph.13126

Chery, J. G., Glos, R. A. E., & Anderson, C. T. (2022). Do woody vines use gelatinous fibers to climb? New Phytologist, 233(1), 126–131. 10.1111/nph.17576

Cho, S. K., Kim, J. E., Park, J.-A., Eom, T. J., & Kim, W. T. (2006). Constitutive expression of abiotic stress-inducible hot pepper *CaXTH3*, which encodes a xyloglucan endotransglucosylase/hydrolase homolog, improves drought and salt tolerance in transgenic *Arabidopsis* plants. FEBS Letters, 580(13), 3136–3144. 10.1016/j.febslet.2006.04.062

Choe, S., Dilkes, B. P., Fujioka, S., Takatsuto, S., Sakurai, A., & Feldmann, K. A. (1998). The DWF4 gene of Arabidopsis encodes a cytochrome P450 that mediates multiple 22alpha-hydroxylation steps in brassinosteroid biosynthesis. The Plant Cell, 10(2), 231–243. 10.1105/tpc.10.2.231

Choi, J. Y., Seo, Y. S., Kim, S. J., Kim, W. T., & Shin, J. S. (2011). Constitutive expression of CaXTH3, a hot pepper xyloglucan endotransglucosylase/hydrolase, enhanced tolerance to salt and drought stresses without phenotypic defects in tomato plants (Solanum lycopersicum cv. Dotaerang). Plant Cell Reports, 30(5), 867–877. 10.1007/s00299-010-0989-3

Clair, B., Alméras, T., Pilate, G., Jullien, D., Sugiyama, J., & Riekel, C. (2011). Maturation Stress Generation in Poplar Tension Wood Studied by Synchrotron Radiation Microdiffraction. Plant Physiology, 155(1), 562–570. 10.1104/pp.110.167270

Clair, B., Déjardin, A., Pilate, G., & Alméras, T. (2018). Is the G-Layer a Tertiary Cell Wall? Frontiers in Plant Science, 9. 10.3389/fpls.2018.00623

Donaldson, L. A., & Singh, A. P. (2016). Chapter 6—Reaction Wood. In Y. S. Kim, R. Funada, & A. P. Singh (Eds.), Secondary Xylem Biology (pp. 93–110). Academic Press. 10.1016/B978-0-12-802185-9.00006-1

Du, J., Gerttula, S., Li, Z., Zhao, S.-T., Liu, Y.-L., Liu, Y., Lu, M.-Z., & Groover, A. T. (2020). Brassinosteroid regulation of wood formation in poplar. New Phytologist, 225(4), 1516–1530. 10.1111/nph.15936

Elmore, C. D., Heatherly, L. G., & Wesley, R. A. (1989). Perennial vine competition and control. Bulletins, 628. https://scholarsjunction.msstate.edu/mafes-bulletins/628

Ferrándiz, C., Liljegren, S. J., & Yanofsky, M. F. (2000). Negative regulation of the SHATTERPROOF genes by FRUITFULL during Arabidopsis fruit development. *Science (New York*, N.Y.), 289(5478), 436–438. 10.1126/science.289.5478.436

Frahm, M. A., Rosas, J. C., Mayek-Pérez, N., López-Salinas, E., Acosta-Gallegos, J. A., & Kelly, J. D. (2004). Breeding beans for resistance to terminal drought in the Lowland tropics. Euphytica, 136(2), 223–232. 10.1023/B:euph.0000030671.03694.bb

Fu, X., Song, A., Peng, B., Li, S., Liu, W., Zhang, L., Jiang, J., Chen, S., & Chen, F. (2023). Brassinosteroid Biosynthetic Gene. Phyton, 92(6), 1681–1694. 10.32604/phyton.2023.027870

Gao, J., Yu, M., Zhu, S., Zhou, L., & Liu, S. (2019). Effects of exogenous 24-epibrassinolide and brassinazole on negative gravitropism and tension wood formation in hybrid poplar (Populus deltoids × Populus nigra). Planta, 249(5), 1449–1463. 10.1007/s00425-018-03074-2

Gartner, B. (1991). Structural stability and architecture of vines vs. Shrubs of poison oak, Toxicodendron diversilobum. Ecology, 72(6), 2005–2015. 10.2307/1941555

Gianoli, E. (2004). Evolution of a climbing habit promotes diversification in flowering plants. Proceedings of the Royal Society B: Biological Sciences, 271(1552), 2011–2015. 10.1098/rspb.2004.2827

Goodstein, D. M., Shu, S., Howson, R., Neupane, R., Hayes, R. D., Fazo, J., Mitros, T., Dirks, W., Hellsten, U., Putnam, N., & Rokhsar, D. S. (2012). Phytozome: A comparative platform for green plant genomics. Nucleic Acids Research, 40(Database issue), D1178–1186. 10.1093/nar/gkr944

Gorshkova, T., Chernova, T., Mokshina, N., Ageeva, M., & Mikshina, P. (2018). Plant ‘muscles’: Fibers with a tertiary cell wall. New Phytologist, 218(1), 66–72. 10.1111/nph.14997

Groover, A. (2016). Gravitropisms and reaction woods of forest trees – evolution, functions and mechanisms. New Phytologist, 211(3), 790–802. 10.1111/nph.13968

Guedes, F. T. P., Laurans, F., Quemener, B., Assor, C., Lainé-Prade, V., Boizot, N., Vigouroux, J., Lesage-Descauses, M.-C., Leplé, J.-C., Déjardin, A., & Pilate, G. (2017). Non-cellulosic polysaccharide distribution during G-layer formation in poplar tension wood fibers: Abundance of rhamnogalacturonan I and arabinogalactan proteins but no evidence of xyloglucan. Planta, 246(5), 857–878. 10.1007/s00425-017-2737-1

Jacobs, J. (2007). *Ecology and Management of field bindweed (Convolvulus arvensis L.)* (Invasive Species Technical Note No. MT-9; Natural Resources Conservation Service). United States Department of Agriculture. https://www.nrcs.usda.gov/plantmaterials/mtpmctn13106.pdf

Ji, S.-J. (2003). Isolation and analyses of genes preferentially expressed during early cotton fiber development by subtractive PCR and cDNA array. Nucleic Acids Research, 31(10), 2534–2543. 10.1093/nar/gkg358

Jourez, B., Riboux, A., & Leclercq, A. (2001). ANATOMICAL CHARACTERISTICS OF TENSION WOOD AND OPPOSITE WOOD IN YOUNG INCLINED STEMS OF POPLAR (POPULUS EURAMERICANA CV ʻGHOY’). IAWA Journal, 22.

Kelly, J. D. (2001). Remaking bean plant architecture for efficient production. In Advances in Agronomy (Vol. 71, pp. 109–143). Elsevier. 10.1016/S0065-2113(01)71013-9

Kim, J. K., & Daniel, G. (2019). Localization of xyloglucan epitopes in the gelatinous layer of developing and mature gelatinous fibers of European aspen (Populus tremula L.) tension wood. BioResources, 14(4), 7675–7686. 10.15376/biores.14.4.7675-7686

Lafarguette, F., Leplé, J.-C., Déjardin, A., Laurans, F., Costa, G., Lesage-Descauses, M.-C., & Pilate, G. (2004). Poplar Genes Encoding Fasciclin-Like Arabinogalactan Proteins Are Highly Expressed in Tension Wood. The New Phytologist, 164(1), 107–121.

Larsson, A. (2014). AliView: A fast and lightweight alignment viewer and editor for large datasets. Bioinformatics, 30(22), 3276–3278. 10.1093/bioinformatics/btu531

Lindgren, C. J., Castro, K. L., Coiner, H. A., Nurse, R. E., & Darbyshire, S. J. (2013). The Biology of Invasive Alien Plants in Canada. 12. Pueraria montana var. Lobata (Willd.) Sanjappa & Predeep. Canadian Journal of Plant Science, 93(1), 71–95. 10.4141/cjps2012-128

Luizon Dias Leme, C., Pace, M. R., & Angyalossy, V. (2021). The “Lianescent Vascular Syndrome” statistically supported in a comparative study of trees and lianas of Fabaceae subfamily Papilionoideae. Botanical Journal of the Linnean Society, 197(1), 25–34. 10.1093/botlinnean/boab015

Mellerowicz, E. J., & Gorshkova, T. A. (2012). Tensional stress generation in gelatinous fibres: A review and possible mechanism based on cell-wall structure and composition. Journal of Experimental Botany, 63(2), 551–565. 10.1093/jxb/err339

Mellerowicz, E. J., Immerzeel, P., & Hayashi, T. (2008). Xyloglucan: The Molecular Muscle of Trees. Annals of Botany, 102(5), 659–665. 10.1093/aob/mcn170

Miller, M. A., Pfeiffer, W., & Schwartz, T. (2010). Creating the CIPRES Science Gateway for inference of large phylogenetic trees. 2010 Gateway Computing Environments Workshop (GCE), 1–8. 10.1109/GCE.2010.5676129

Nishikubo, N., Awano, T., Banasiak, A., Bourquin, V., Ibatullin, F., Funada, R., Brumer, H., Teeri, T. T., Hayashi, T., Sundberg, B., & Mellerowicz, E. J. (2007). Xyloglucan Endo-transglycosylase (XET) Functions in Gelatinous Layers of Tension Wood Fibers in Poplar—A Glimpse into the Mechanism of the Balancing Act of Trees. Plant and Cell Physiology, 48(6), 843–855. 10.1093/pcp/pcm055

Onyenedum, J. G., Sousa-Baena, M. S., Hunt, L. M., Acevedo, A. A., Glos, R. A. E., & Anderson, C. T. (2025). Gelatinous fibers develop asymmetrically to support bends and coils in common bean vines (Phaseolus vulgaris). American Journal of Botany, 112(3), e70014. 10.1002/ajb2.70014

Paul, G. S., & Yavitt, J. B. (2011). Tropical Vine Growth and the Effects on Forest Succession: A Review of the Ecology and Management of Tropical Climbing Plants. The Botanical Review, 77(1), 11–30. 10.1007/s12229-010-9059-3

Phaseolus vulgaris 2.1. (2020). Phytozome, Joint Genome Institute. https://data.jgi.doe.gov/refine-download/phytozome?genome_id=442&_gl=1*pobr41*_ga*NTc1NDAzNzYwLjE3NTQ5MzYzMDM.*_ga_YBLMHYR3C2*czE3NTQ5MzYzMDIkbzEkZzAkdDE3NTQ5MzYzMDIkajYwJGwwJGgw

R Core Team. (2025). *R: A Language and Environment for Statistical Computing* (Version 4.4.3) [Computer software]. R foundation for Statistical Computing. https://www.R-project.org/

Rambaut, A., & Drummond, A. J. (2012). *Rambaut: FigTree version 1.4.0* [Computer software].

Ricklefs, R. E., & Renner, S. S. (1994). Species Richness Within Families of Flowering Plants. Evolution, 48(5), 1619–1636. 10.1111/j.1558-5646.1994.tb02200.x

Sandler, H. A., & Ghantous, K. (2019). Dodder: Biology and Management. Cranberry Station Fact Sheets, 43. https://hdl.handle.net/20.500.14394/9127

Scheller, H. V., & Ulvskov, P. (2010). Hemicelluloses. Annual Review of Plant Biology, 61(Volume 61, 2010), 263–289. 10.1146/annurev-arplant-042809-112315

Schindelin, J., Arganda-Carreras, I., Frise, E., Kaynig, V., Longair, M., Pietzsch, T., Preibisch, S., Rueden, C., Saalfeld, S., Schmid, B., Tinevez, J.-Y., White, D. J., Hartenstein, V., Eliceiri, K., Tomancak, P., & Cardona, A. (2012). Fiji: An open-source platform for biological-image analysis. Nature Methods, 9(7), Article 7. 10.1038/nmeth.2019

Schnitzer, S. A. (2024). The Ecology of Lianas and Their Increasing Influence in Tropical Forests. In Routledge Handbook of Forest Ecology (2nd ed.). Routledge.

Sivan, P., Mishra, P., & Rao, K. S. (2010). Occurrence of Reaction xylem in the peduncle of Couroupita Guianensis and Kigelia Pinnata. 10.1163/22941932-90000017

Song, L., Valliyodan, B., Prince, S., Wan, J., & Nguyen, H. T. (2018). Characterization of the XTH Gene Family: New Insight to the Roles in Soybean Flooding Tolerance. International Journal of Molecular Sciences, 19(9), Article 9. 10.3390/ijms19092705

Sousa-Baena, M. S., & Onyenedum, J. G. (2022). Bouncing back stronger: Diversity, structure, and molecular regulation of gelatinous fiber development. Current Opinion in Plant Biology, 67, 102198. 10.1016/j.pbi.2022.102198

Sousa-Baena, M. S., Sinha, N. R., Hernandes-Lopes, J., & Lohmann, L. G. (2018). Convergent Evolution and the Diverse Ontogenetic Origins of Tendrils in Angiosperms. Frontiers in Plant Science, 9. 10.3389/fpls.2018.00403

Sperotto, P., Acevedo-Rodríguez, P., Vasconcelos, T. N. C., & Roque, N. (2020). Towards a Standardization of Terminology of the Climbing Habit in Plants. The Botanical Review, 86(3), 180–210. 10.1007/s12229-020-09218-y

Sperotto, P., Roque, N., Acevedo-Rodríguez, P., & Vasconcelos, T. (2023). Climbing mechanisms and the diversification of neotropical climbing plants across time and space. New Phytologist, 240(4), 1561–1573. 10.1111/nph.19093

Stamatakis, A. (2014). RAxML version 8: A tool for phylogenetic analysis and post-analysis of large phylogenies. Bioinformatics, 30(9), 1312–1313. 10.1093/bioinformatics/btu033

Stoffers, A. L. (1979). Fabaceae. Flora of the Netherlands Antilles, 3(2), 61–123.

Sugiyama, K., Okuyama, T., Yamamoto, H., & Yoshida, M. (1993). Generation process of growth stresses in cell walls: Relation between longitudinal released strain and chemical composition. Wood Science and Technology, 27(4), 257–262. 10.1007/BF00195301

Symons, G. M., & Reid, J. B. (2004). Brassinosteroids Do Not Undergo Long-Distance Transport in Pea. Implications for the Regulation of Endogenous Brassinosteroid Levels. Plant Physiology, 135(4), 2196–2206. 10.1104/pp.104.043034

Szekeres, M., Németh, K., Koncz-Kálmán, Z., Mathur, J., Kauschmann, A., Altmann, T., Rédei, G. P., Nagy, F., Schell, J., & Koncz, C. (1996). Brassinosteroids Rescue the Deficiency of CYP90, a Cytochrome P450, Controlling Cell Elongation and De-etiolation in Arabidopsis. Cell, 85(2), 171–182. 10.1016/S0092-8674(00)81094-6

Takahashi, D., Johnson, K. L., Hao, P., Tuong, T., Erban, A., Sampathkumar, A., Bacic, A., Livingston III, D. P., Kopka, J., Kuroha, T., Yokoyama, R., Nishitani, K., Zuther, E., & Hincha, D. K. (2021). Cell wall modification by the xyloglucan endotransglucosylase/hydrolase XTH19 influences freezing tolerance after cold and sub-zero acclimation. Plant, Cell & Environment, 44(3), 915–930. 10.1111/pce.13953

Takeuchi, M., Guedes, F. T. P., Laurans, F., Boizot, N., Déjardin, A., & Pilate, G. (2024). Immunolocalization of fasciclin-like arabinogalactan proteins in the G-layers of poplar tension wood fibers (p. 2024.12.19.629400). bioRxiv. 10.1101/2024.12.19.629400

Tomlinson, P. B., Magellan, T. M., & Griffith, M. P. (2014). Root contraction in Cycas and Zamia (Cycadales) determined by gelatinous fibers. American Journal of Botany, 101(8), 1275–1285. 10.3732/ajb.1400170

van der Heijden, G. M., Schnitzer, S. A., Powers, J. S., & Phillips, O. L. (2013). Liana Impacts on Carbon Cycling, Storage and Sequestration in Tropical Forests. Biotropica, 45(6), 682–692. 10.1111/btp.12060

Van Sandt, V. S. T., Suslov, D., Verbelen, J.-P., & Vissenberg, K. (2007). Xyloglucan Endotransglucosylase Activity Loosens a Plant Cell Wall. Annals of Botany, 100(7), 1467–1473. 10.1093/aob/mcm248

Vaughn, K. C., & Bowling, A. J. (2011). Biology and Physiology of Vines. In Horticultural Reviews (Vol. 38). Wiley-Blackwell. https://www.ars.usda.gov/ARSUserFiles/60663500/Publications/Vaughn/Vaughn%20et%20al._2011_Biology%20and%20physiology%20of%20vines.pdf

Vukašinović, N., Wang, Y., Vanhoutte, I., Fendrych, M., Guo, B., Kvasnica, M., Jiroutová, P., Oklestkova, J., Strnad, M., & Russinova, E. (2021). Local brassinosteroid biosynthesis enables optimal root growth. Nature Plants, 7(5), 619–632. 10.1038/s41477-021-00917-x

Wang, H., Jin, Y., Wang, C., Li, B., Jiang, C., Sun, Z., Zhang, Z., Kong, F., & Zhang, H. (2017). Fasciclin-like arabinogalactan proteins, PtFLAs, play important roles in GA-mediated tension wood formation in Populus. Scientific Reports, 7(1), 6182. 10.1038/s41598-017-06473-9

White, D. J. B., & Robards, A. W. (1965). Gelatinous Fibres in Ash (Fraxinus excelsior L.). Nature, 205(4973), 818–818. 10.1038/205818a0

Wyka, T. P., Oleksyn, J., Karolewski, P., & Schnitzer, S. A. (2013). Phenotypic correlates of the lianescent growth form: A review. Annals of Botany, 112(9), 1667–1681. 10.1093/aob/mct236

Xuan, Y., Zhou, Z. S., Li, H. B., & Yang, Z. M. (2016). Identification of a group of XTHs genes responding to heavy metal mercury, salinity and drought stresses in *Medicago truncatula*. Ecotoxicology and Environmental Safety, 132, 153–163. 10.1016/j.ecoenv.2016.06.007

Yoshimitsu, Y., Tanaka, K., Fukuda, W., Asami, T., Yoshida, S., Hayashi, K.-I., Kamiya, Y., Jikumaru, Y., Shigeta, T., Nakamura, Y., Matsuo, T., & Okamoto, S. (2011). Transcription of DWARF4 plays a crucial role in auxin-regulated root elongation in addition to brassinosteroid homeostasis in Arabidopsis thaliana. PloS One, 6(8), e23851. 10.1371/journal.pone.0023851

Zebosi, B., Vollbrecht, E., & Best, N. B. (2024). Brassinosteroid biosynthesis and signaling: Conserved and diversified functions of core genes across multiple plant species. Plant Communications, 5(9). 10.1016/j.xplc.2024.100982

Zimmermann, M. H., Wardrop, A. B., & Tomlinson, P. B. (1968). Tension wood in aerial roots of ficus benjamina L. Wood Science and Technology, 2(2), 95–104. 10.1007/BF00394958

